# A novel luminescence-based β-arrestin membrane recruitment assay for unmodified GPCRs

**DOI:** 10.1101/2020.04.09.034520

**Authors:** Maria Hauge Pedersen, Jennifer Pham, Helena Mancebo, Asuka Inoue, Jonathan A. Javitch

**Author notes:** Corresponding author: Jonathan A. Javitch.

## Abstract

G protein-coupled receptors (GPCRs) signal through activation of G proteins and subsequent modulation of downstream effectors. More recently, G protein-independent signaling via the arrestin pathway has also been implicated in important physiological functions. This has led to great interest in the identification of biased ligands that favor either the G protein or arrestin-signaling pathways. Currently available screening techniques that measure arrestin recruitment have required C-terminal receptor modifications that can in principle alter protein interactions and thus signaling. Here, we have developed a novel luminescence-based assay to measure arrestin recruitment to any unmodified receptor.

NanoLuc, an engineered luciferase from *ophlorus gracilirostris* (deep sea shrimp), is smaller and brighter than other well-established luciferases. Recently, several publications have explored functional NanoLuc split sites for use in complementation assays. Here, we have identified a novel split site and have fused the N-terminal fragment to a membrane tether and the C-terminal fragment to the N-terminus of either β-arrestin 1 or 2. Upon receptor activation, arrestin is recruited to the plasma membrane in an agonist concentration-dependent manner and the two NanoLuc fragments complement to reconstitute functional luciferase, which allows quantification of recruitment with a single luminescence signal. Our assay avoids potential artifacts related to C-terminal receptor modification. The split NanoLuc arrestin recruitment assay has promise as a new generic assay for measuring arrestin recruitment to diverse GPCR types in heterologous or native cells.

## Introduction

Over the past decade, increasing attention has been directed to biased signaling of GPCRs and the possibility of identifying ligands that can selectively target either G protein-mediated signaling or G protein-independent signaling through β-arrestin. As an example, a “β-blocker” that antagonizes G protein activation but recruits β-arrestin has been found to increase survival rates in patients suffering from heart failure (1). At the μ-opioid receptor G protein signaling has been proposed to be responsible for analgesia, with side effects such as respiratory depression resulting from arrestin-mediated signaling (2,3), although this paradigm has recently been challenged (4–6). We have demonstrated that β-arrestin recruitment to the dopamine D2 receptor in indirect pathway neurons in the ventral striatum leads to enhanced locomotion, whereas G protein signaling is necessary for incentive behavior (7), further emphasizing the potential importance of biased signaling in more targeted therapeutics.

The most well-established techniques to investigate β-arrestin recruitment are the PathHunter, Tango, LinkLight, and Bioluminescence resonance energy transfer (BRET) assays, all of which require C-terminal modifications of the receptor of interest. The PathHunter assay is based on an enzyme fragment complementation assay in which the split enzyme fragments are attached to the receptor and to β-arrestin and complementation of the functional enzyme creates a chemiluminescence readout when substrate is added (8). The Tango GPCR assay is a reporter gene assay in which a transcription factor attached to the receptor is cleaved off by a protease-tagged β-arrestin, which leads to the expression of a reporter that creates a luminescence readout upon substrate addition (9,10). LinkLight uses a modified luciferase attached to β-arrestin that is cleaved off by a protease attached to the GPCR of interest, thereby activating the luciferase and producing light (11). BRET between a receptor fused at its C-terminus to a luciferase and arrestin tagged at its N-terminus with mVenus is also commonly used to measure arrestin recruitment (12). Notably, all of these β-arrestin assays require C-terminal fusions to the receptor of interest.

In contrast, we previously introduced a modified BRET assay in which the donor (Rluc8) is anchored to the membrane and the acceptor (Venus) is fused to β-arrestin (12). In this assay, which is based on “bystander” BRET, translocation of β-arrestin to the receptor at the plasma membrane leads to an increase in BRET from a membrane-anchored donor. This assay relies on yeast-derived “helper” peptides that are attached to the donor and acceptor, thereby enhancing the affinity of the interaction between arrestin and the plasma-membrane anchor (13). This bystander BRET format was later adapted to a BRET2 format (14). Both these assays require a dual output microplate reader compatible with BRET. The novel luciferase NanoLuc developed by Promega has the advantages of being smaller and brighter compared to the other known luciferases (15). Recently, split NanoLuc, known as NanoBiT, has been used successfully in a direct recruitment assay in which the small “bit” is attached to the receptor of interest and the large “bit” is attached to β-arrestin. We expected that NanoBiT could readily be adapted to our bystander assay by attaching our membrane-anchor to the small bit instead of the receptor, thereby duplicating our membrane recruitment assay with complemented NanoLuc. To our surprise, these split constructs failed when adapted to the membrane-recruitment assay, as we detected no increase in luminescence with activation of the dopamine D2, β2 adrenergic or angiotensin AT1 receptors (D2R, β2AR, AT1R, respectively).

Here we have developed a novel split site for NanoLuc that is suitable for the desired NanoLuc-complementation-based β-arrestin membrane recruitment assay in which the <u>Me</u>mbrane anchor is attached to the <u>N</u>-terminal fragment (referred to as MeN) and β-<u>Arr</u>estin is attached to the <u>C</u>-terminal fragment (referred to as ArC), creating an assay that does not rely on receptor modification or helper peptides and provides a simple luminescence output reading of β-arrestin recruitment to investigate receptors in a more native format. The novel assay named MeNArC worked for all tested receptors and can serve as a general assay for β-arrestin recruitment to unmodified GPCRs both *in vitro* and potentially *in vivo*.

## Results

### Novel NanoLuc split site forms a functional protein upon complementation and works in direct arrestin-receptor recruitment

Given the failure of the commercial NanoBiT complementation to detect agonist-induced arrestin recruitment when adapted to our desired membrane recruitment format (Fig. S1), we chose a new split site based on the crystal structure of NanoLuc (16). We selected a site in a loop region that divides NanoLuc into two almost equally sized fragments (amino acids 1-102 and 103-172) without disrupting any secondary structural elements, in the hope that the fragments would fold independently and efficiently reassemble when brought together at the membrane (Fig. 1A). This site was similar but not identical to a Nanoluc split site used previously in a direct recruitment assay (17). To verify the ability of our split fragments to form a fully functional protein when complemented, we first attached the N- and C-terminal NanoLuc fragments to FRB (FKBP-rapamycin binding domain) and FKBP (FK506- and rapamycin binding protein), respectively, which can readily dimerize with high affinity upon addition of rapamycin (Fig. 1B)(18). When transfected alone, the fragments did not yield any luminescence over the baseline seen with mock transfected cells (Fig. 1C).

**Figure 1.**
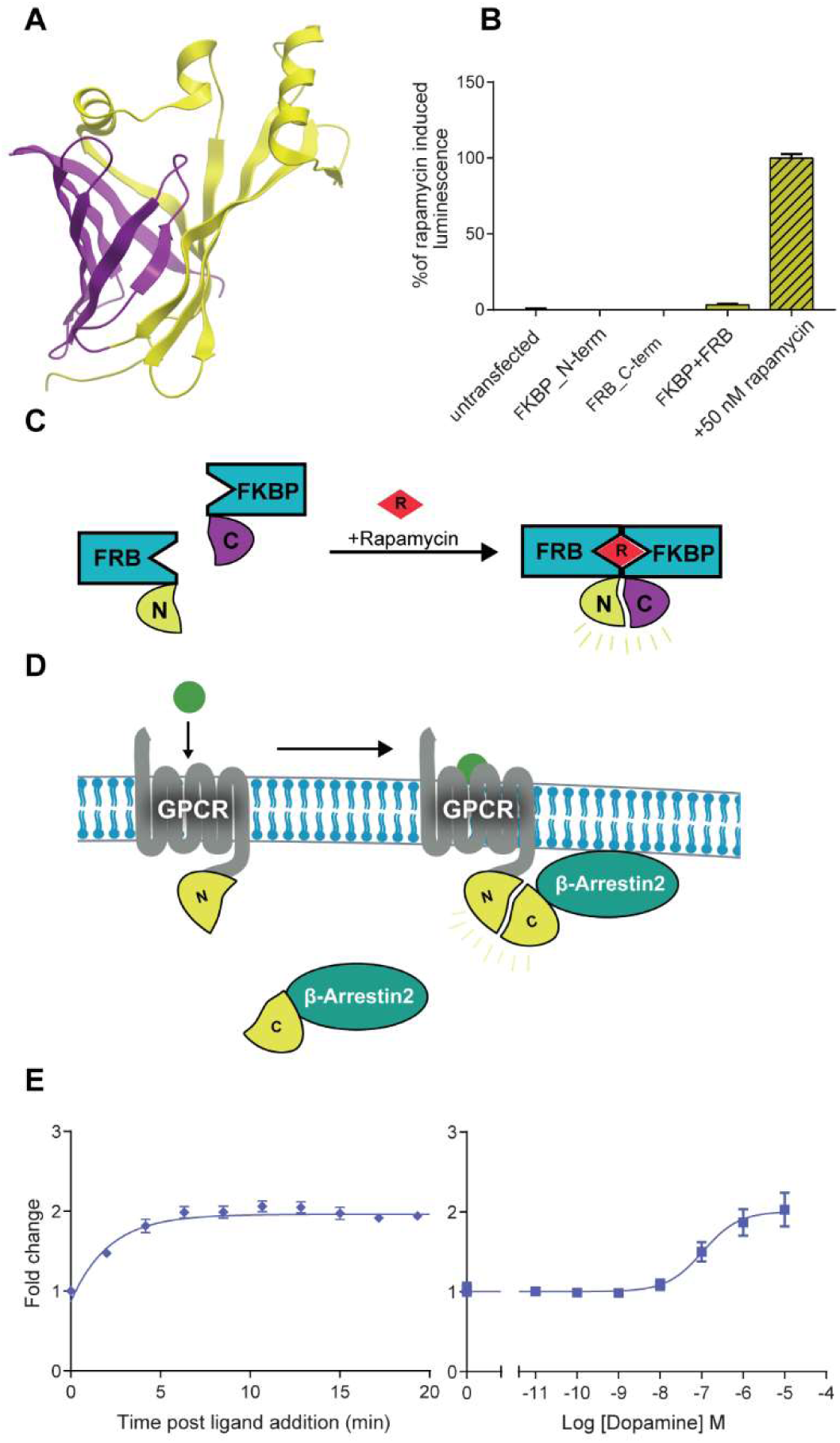
A) NanoLuc split visualized with Molsoft Browser® showing the N- and C-terminal components, in yellow and purple, respectively (PDB #5IBO) C) Schematic overview of rapamycin-induced dimerization of FRB and FKBP with fused N- and C-terminal NanoLuc splits respectively. B) Hek293 cells were transiently transfected with either FRB_N-term, FKBP_C-term or both +/- 50 nM rapamycin overnight. 50 µM Coelenterazine H was added and luminescence detected. Single transfections elicited no luminescence. When constructs were cotransfected, a low degree of self-assembly yielded 3% of the rapamycin-induced complementation-based luminescence. D) Schematic overview of direct recruitment assay, where the N-terminal split is fused to the C terminus of the receptor and the C-terminal split is fused to the N terminus of β-arrestin2. E) Direct recruitment assay with D2R. Time course after dopamine addition on the left and dose response curve on the right. All data represent mean ± SEM of 3-5 independent experiments with triplicate determinations.

Cotransfection of both fragments yielded a small basal luminescence signal that was increased ∼30-fold by the addition of rapamycin, suggesting only minimal spontaneous assembly of the fragments and efficient complementation when brought into proximity, demonstrating that the new fragments are suitable as a tool for complementation assays.

Next, we tested the split fragments in direct β-arrestin recruitment with a well-established but weak arrestin recruiter (19,20), the dopamine D2 receptor (D2R), by attaching the N-terminal NanoLuc fragment to the C terminus of D2R and the C-terminal NanoLuc fragment to the N terminus of β-arrestin2 (Fig. 1D). Upon addition of dopamine, luminescence increased, reaching a plateau after ∼6 min with a 2-fold increase (EC50 = 108 nM± 20) (Fig. 1E), comparable to DiscoverX’s pathhunter assay (EC50 = 90 nM± 30) and BRET direct recruitment assay (EC50 = 50 nM± 10) (21). Having shown the utility of the split fragments in a complementation assay reading out on the direct recruitment of β-arrestin2 to the receptor, we moved forward to see if we could use the splits in a β-arrestin membrane recruitment assay with unmodified receptors.

### Establishing the utility of the MeNArC assay in the bystander configuration

We modified our previously developed β-arrestin membrane recruitment BRET assay (12) by exchanging the donor and acceptor with our NanoLuc fragments, with the N-terminal fragment attached to the membrane (double palmitoylated fragment of GAP43) and the C-terminal part attached to the N terminus of β-arrestin2 (Fig. 2A).

**Figure 2.**
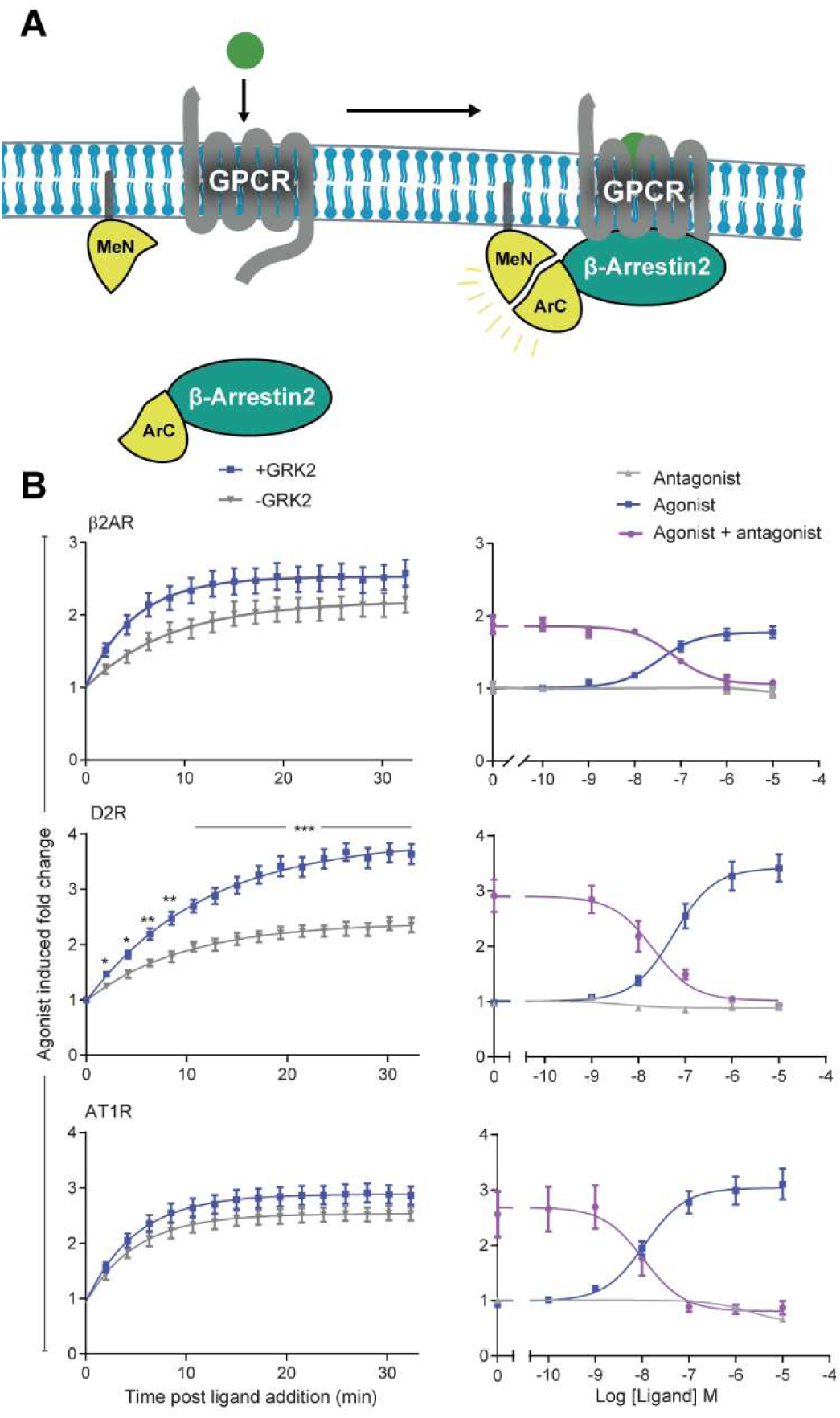
A) Schematic overview of the arrestin membrane-recruitment assay. The N-terminal fragment is fused to a doubly palmitoylated fragment of GAP43, thereby tethering it to the membrane (MeN) and the C-terminal fragment is fused to the N terminus of β-arrestin2 (ArC). B) Arrestin recruitment shown in fold change of luminescence with 3 different receptors transiently transfected in Hek293 cells; Angiotensin 1 receptor (AT1R), Beta2 adrenergic receptor (β2AR) and the Dopamine 2 receptor (D2R) respectively with time course for agonist-induced arrestin recruitment in the presence (blue) or absence (grey) of GRK2 on the left panels. Steady state is reached after 20 min for all receptors. Right panels show dose-response curves of arrestin recruitment with agonist (blue line), antagonist (grey line) and a high dose of agonist and antagonist (purple line) cotransfected with GRK2. Agonist/ antagonist used for the respective receptors were angiotensin II/olmesartan (AT1R), isoproterenol/propranolol (β2AR) and dopamine/sulpiride (D2R). pEC50 for β2AR, D2R & AT1R was −7.50±0.14, −7.25±0.16 & −7.94±0.14 respectively.

To investigate the versatility of the MeNArC assay we tested it with the β2 adrenergic receptor (β2AR) and dopamine D2R, as representative class A receptors, and angiotensin 1 receptor (AT1R), a class B receptor. This classification is based on their agonist-induced β-arrestin binding profile after internalization, with class A receptors binding β-arrestin more transiently compared to class B receptors, which bind with higher affinity and form a longer-lived stable complex (22) (Fig. 2C). Agonist-induced β-arrestin recruitment, as measured by an increase in luminescence, reached a plateau within ∼20 min for all three receptors. (Fig. 2C left panel). Consistent with previous findings (23), co-transfection with GRK2 substantially increased β-arrestin recruitment for D2R with a fold change of: 3.9±0.15/2.4±0.08 (+/-GRK2, respectively) but only had a minor effect on β2AR 2.53±0.07/2.19±0.11, +/-GRK2) and AT1R (2.89±0.07/2.53±0.05, +/-GRK2).

We next tested if the interaction was reversible by adding antagonist 20 min after agonist addition. Antagonist addition with either β2AR and D2R led to a strong inhibition of luminescence down to baseline (Fig. 2B right panel), showing that the complementation is reversible and can therefore also be used to investigate inverse agonism in a receptor with sufficient constitutive activity. In contrast, whereas the AT1R antagonist olmesartan prevented the agonist-induced increase if added prior to agonist, it failed to decrease luminescence when added after agonist (data not shown). We reasoned that this reflects the much higher affinity of β-arrestin interaction with AT1R once the C tail is phosphorylated (22). Like for the β2AR-V2R chimera, we infer that even after blocking core engagement of AT1R with arrestin by addition of the antagonist, the interaction with the phosphorylated C tail is sufficient to maintain the interaction (24), in contrast to the β2AR and D2R where core engagement and therefore continued agonist binding are essential for arrestin interaction.

We also applied the MeNArC assay to the µ-opioid receptor and cannabinoid 1 receptor, two other class A receptors, with similar results (Fig. S2). We also tested the assay with β-arrestin1 and D2R with and without coexpression of GRK2. This assay format was also successful, but with a slightly lower dynamic range (fold change: 3.1±0.14/1.92± 0.14, +/-GRK2). When tested in antagonist mode with D2R and β-arrestin1(Fig. S3) we found the luminescence to be inhibited, similar to the findings with β-arrestin2 (Fig. 2).

### Creation of a multicistronic vector system to use MeNArC in stably transfected CHO cell lines

For use in *in vitro* and potentially *in vivo*, we designed a multicistronic vector for MeNArC where the 2 constructs expressing the MeNArC components were inserted into one vector separated by the self-cleaving peptide P2A (25). As a test case, cell lines stably expressing each of the opioid receptors were stably transfected with the multicistronic MeNArC vector and individual clones were chosen for intermediate luminescence level (see Methods). µ, δ, κ and nociceptin opioid receptors (MOR, DOR, KOR and NOP respectively) all exhibited robust dose-response curves with their respective agonists (Fig. 3) yielding fold changes of MOR: 3.32±0.05; KOR: 1.72±0.06; DOR: 2.46±0.13 and NOP: 1.50±0.03, all with unmodified receptors in the absence of exogenous GRK2.

**Figure 3.**
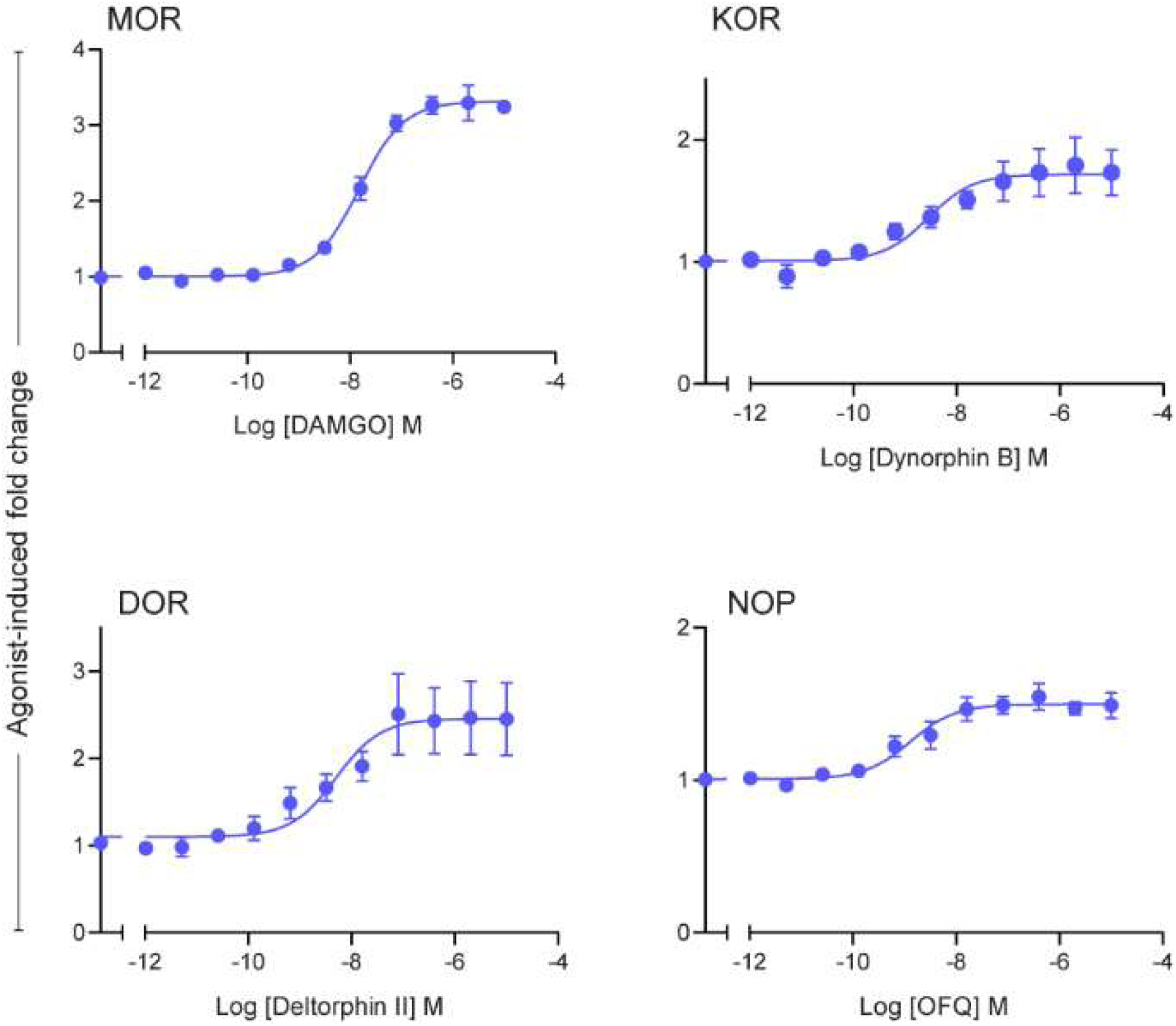
Dose response curves of arrestin recruitment for each of the opioid receptors; MOR (μ), KOR (κ), DOR (δ) or NOP (nociception), stably transfected into Cho-k1 cells together with the β-arrestin2 version of the ArC-P2A-MeN plasmid.

## Discussion

We have developed a novel NanoLuc complementation assay that measures β-arrestin recruitment to non-modified receptors using a simple luminescence output. The most widely used β-arrestin assays available all rely on modified receptors, which can have unwanted effects on the outcome, as part of the reporter is fused to the C-terminal tail of the receptor that is also the primary recognition site for β-arrestins in most receptors, which might alter receptor-arrestin interactions (26). The only other available assays, which also use unmodified receptors, are the transfluor assay where the reading output is microscopy-based and the aforementioned BRET method. which require more specialized equipment.

Complementation-based luciferase split assays have been widely used in the last decade to detect protein-protein interaction in multiple biological systems; NanoLuc has advantages in both decreased size and increased brightness (15). Several NanoBiT-based GPCR complementation methods have been developed, including direct β-arrestin recruitment, direct G protein interaction and internalization, all of which have the small fragment fused to the C terminus of the receptor (27–29). Curiously, we found that the NanoBiT system, when adapted to the β-arrestin membrane recruitment assay, failed to show a response with any of the three receptors tested, including the D2R (Fig. S1). In contrast, a recent publication showed that NanoBiT was successful for a direct recruitment assay with the D2R fused to the small bit (28). We do not know why these split fragments work with the direct recruitment assay but not the membrane recruitment assay, but we speculated that this might result from unfavorable orientation of the attachments of the large and small fragments. Thus, we tried splitting NanoLuc into two roughly equal sized fragments

We tested our membrane recruitment assay with the opioid receptors, CB1R, D2R, β2AR & AT1R. The MeNArC assay also proved to be reversible for class A receptors, which indicates its potential use for testing inverse agonists. We further adapted the method for stable transfection, generating a single plasmid containing both MeNArC components for the creation of cell lines amenable to high throughput screening. An advantage of this assay is that it can be used, in principle, in biological systems with native receptors both in vitro and potentially in vivo. While our assay read outs on arrestin recruitment, like all arrestin recruitment assays it does not measure β-arrestin activity and therefore does not directly report on downstream signaling, which can involve several different pathways such as ERK and Src activation, as well as internalization (30). Another important note is that even though the membrane recruitment assay can be used with unmodified receptors to give a more natural picture of β-arrestin recruitment upon receptor activation, we cannot rule out action of the tested agonist at endogenous receptors or possible off-target receptors when tested in native tissues unless knockout models are available.

In conclusion, we have here developed a novel β-arrestin membrane recruitment assay using complementation of novel NanoLuc fragments that we believe can serve as a generic and readily adapted screening method for detecting β-arrestin recruitment to unmodified receptors.

## Materials and Methods

### Compounds

Dopamine hydrochloride (#H-8502) and angiotensin II (#A9525) were purchased from Sigma-Aldrich (St. Louis, MO, USA); sulpiride (#0895), olmesartan (#4616), Dynorphin B (#3196), naloxone hydrochloride (#0599), WN552122 (#1038) and rimonabant hydrochloride (#0923) from Tocris (Bristol, UK); DAMGO (#024-10) from Phoenix Pharmaceuticals and rapamycin (#tlrl-rap) from Invivogen (San Diego, CA, USA).

### Plasmid constructs

All constructs were cloned into pcDNA3.1+ (Invitrogen, Carlsbad, CA). All constructs were made using site-directed mutagenesis. NanoLuc was split to create an N-terminal part (N1) with amino acids 1-102 and a C-terminal part (N2) with amino acids 103-172. N1 and N2 were attached with an 8 amino acid flexible linker at the C-terminal end of FKBP and N2 at the C-terminal end of FRB (FKBP rapamycin binding domain of mTOR), both with a signal peptide (MKTIIALSYIFCLVFA) and myc (EQKLISEEDL) tag at the N-terminal of FRB/FKBP. NanoLuc was codon optimized for mammalian expression changing 87 bp (see supplementary Fig. 2 for sequence). Human dopamine 2 receptor, CB1R and β2AR had the signal peptide shown previously and a FLAG (ADYKDDDDA) tag attached to the N-terminal. Mammalian angiotensin receptor 1 (AT1R) had a signal peptide and snap tag attached to the N-terminal. The constructs for the direct and indirect β-arrestin recruitment assays were made by exchanging the donor and acceptor with the NanoLuc splits from the BRET arrestin method previously described (12) as well as removal of the helper peptides SH3 and Sp1 to make D2R-linker-N1, mem-linker-N1 (membrane tethered N-terminal split named MeN) and N2-linker-β-arrestin2 (arrestin tethered C-terminal split named ArC). The multicistronic vector was made by inserting the 22 amino acid P2A cleavage peptide between sequence of MeN and ArC, creating ArC-P2A-MeN in pcDNA3.1+ hygromycin. GRK2 was obtained from David Sibley. The human μ/κ/δ/Nociceptin opioid receptors were all inserted into the pMEX2 vector from Multispan.

To compare Promega’s NanoBiT system β-arrestin2 was inserted at the C-terminal end of LgBiT using Promegas pBiT1.1-N vector (Promega Cat. #N2014) and the doubly palmitolyated fragment of GAP43 was inserted at the N-terminal of SmBiT using the pBiT2.1-C vector.

### Cell culture and transfection

Hek293 cells (ATTC® CRL-1573) were grown in DMEM + GlutaMAX™-I (Gibco, Invitrogen, Paisley, UK) with 10% fetal bovine serum (Cornig #35-010) and 1% penicillin/streptomycin (Cornig #30-002) at 37 °C with 5% CO_2_. Cells were transfected with polyethylenimine (PEI; linear, MW 25,000; Polysciences, cat. No. 23966-2) at a 2:1 ratio (PEI:DNA) in growth medium.

CHO-k1 cell lines stably transfected with either the μ/κ/δ/Nociceptin opioid receptor were further transfected with ArC-P2A-MeN and single clones were chosen with an intermediate luminescence level. The cell lines were grown in HAM F12 with 1% L-glutamine, 10% fetal bovine serum, 10 μg/ml puromycin (invivogen ant-pr-5), 800 µg/ml G418 (Corning 30-234-CI) and 1% penicillin/streptomycin at 37 °C with 5% CO_2_.

### Arrestin complementation assay

Two days after transfection the Hek293 cells were washed with PBS and seeded into a 96 well black-white iso plate (Perkin Elmer) in pbs containing 5 mM glucose and supplemented with 5 µM of Nanoglo (Promega #N1120) or coelenterazine-H (Dalton #50909-86-9). After 5 min incubation at room temperature with a foil cover, ligand was added and luminescence measured every 2 min for kinetics data or after 20 min for drug response curves using a Pherastar FS (BMG labtech) plate reader.

The stably transfected CHO cells were seeded into a white opaque bottom Poly-D-lysine-coated 384 well plate (Corning BioCoat™ #356661). Cells were washed with HBSS and incubated with 5 μM coelenterazine-H for 5 min before compound addition. Luminescence was read 20 min after incubation on a Flexstation III.

## Data analysis

Data were analyzed using GraphPad Prism Version 7.

## Acknowledgements

Supported by NIH grant MH54137 (J.A.J) and NNF grant NNF17OC0024830 (M.H.P.)

---

**Supplementary Figure 1.**
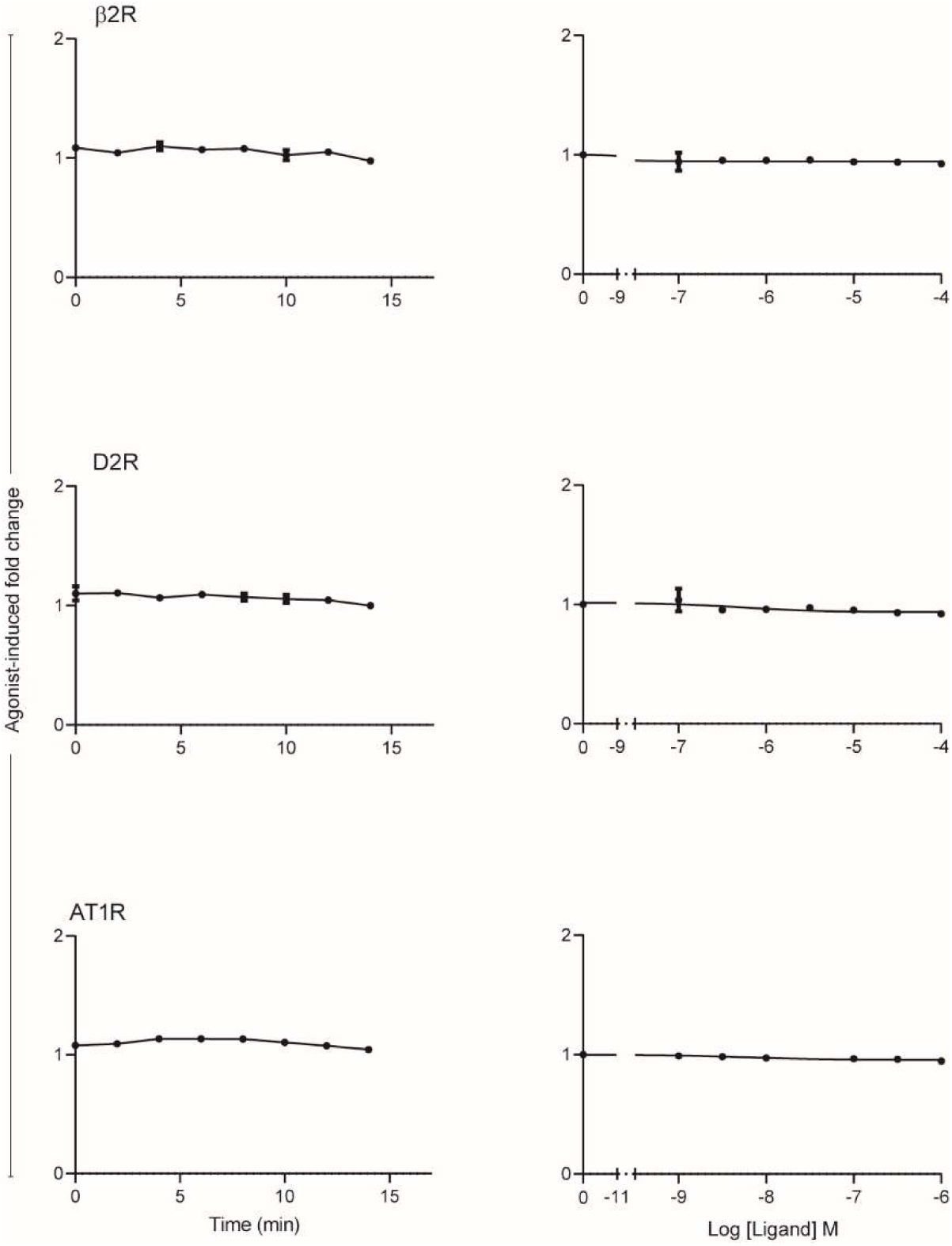
arrestin membrane recruitment assay tested with NanoBiT NanoBiT tested as an β-arrestin2 membrane-recruitment assay where the small N-terminal split SmBiT was tethered to the membrane by fusing it to the doubly palmitolyated fragment of GAP43 and the C-terminal LgBiT was fused to the N terminus of β-arrestin2. Tested with 3 different receptors; Angiotensin 1 (AT1R), beta2-adrenergic receptor (β2AR) and dopamine 2 (D2R). Shown are time course curves after agonist addition on the left and dose response curves on the right, with the agonist isoproterenol, quinpirole and angiotensin II used for β2AR, D2R and AT1R respectively.

**Supplementary Figure 2.**
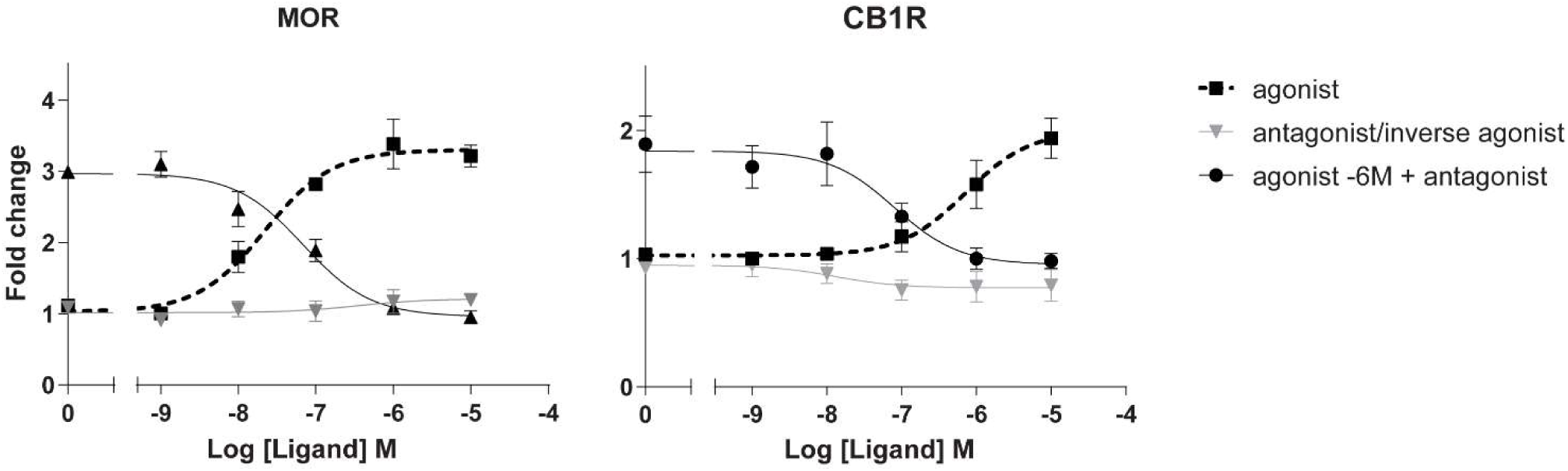
MeNArC assay with dose-response curves for β-arrestin2 membrane recruitment with the µ-opioid receptor (MOR) and cannabinoid 1 receptor (CB1R). The agonist DAMGO and the antagonist naxolone were used for MOR and the agonist WN552122 and antagonist rimonabant were used for CB1R.

**Supplementary Figure 3.**
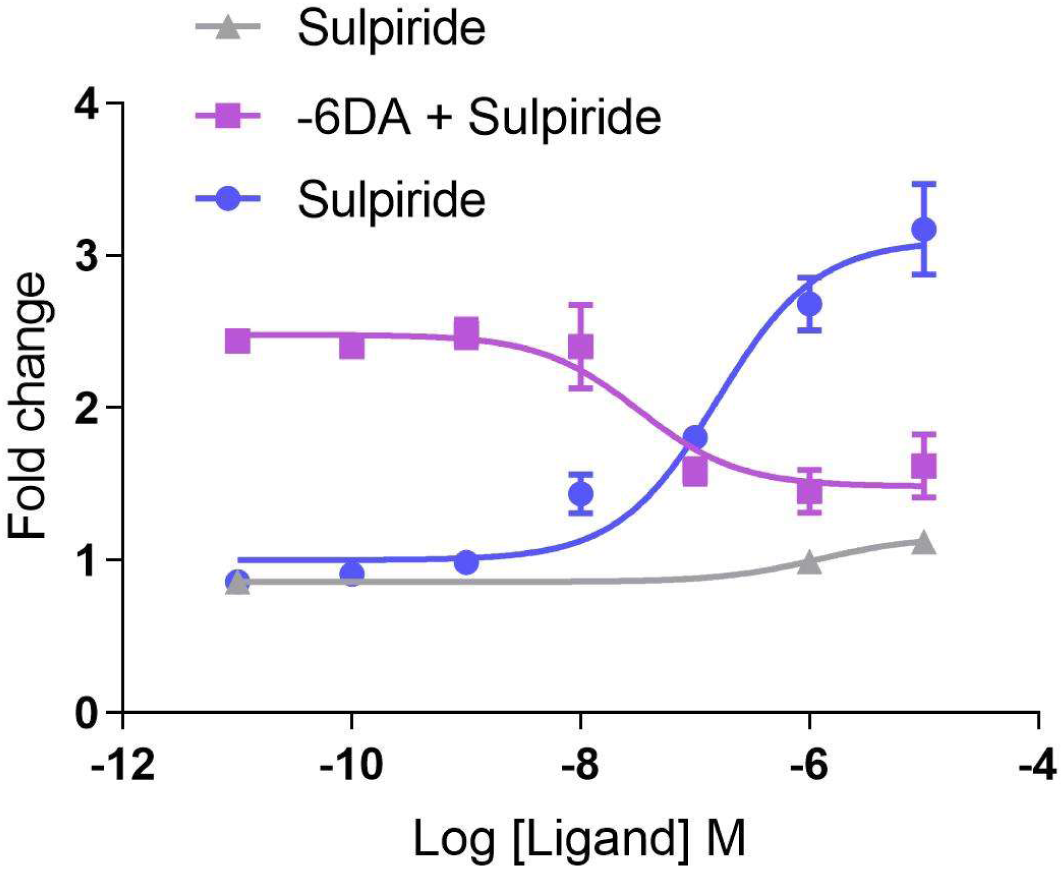
D2R membrane arrestin recruitment with β-arrestin1 Dose response curves of the membrane arrestin recruitment assay tested with β-arrestin1 and D2R cotransfected with GRK2. With the endogenous agonist dopamine in blue, the antagonist sulpiride alone in grey and 10 μM dopamine with the addition of sulpiride 20 min after agonist addition.

**Supplementary Figure 4.**
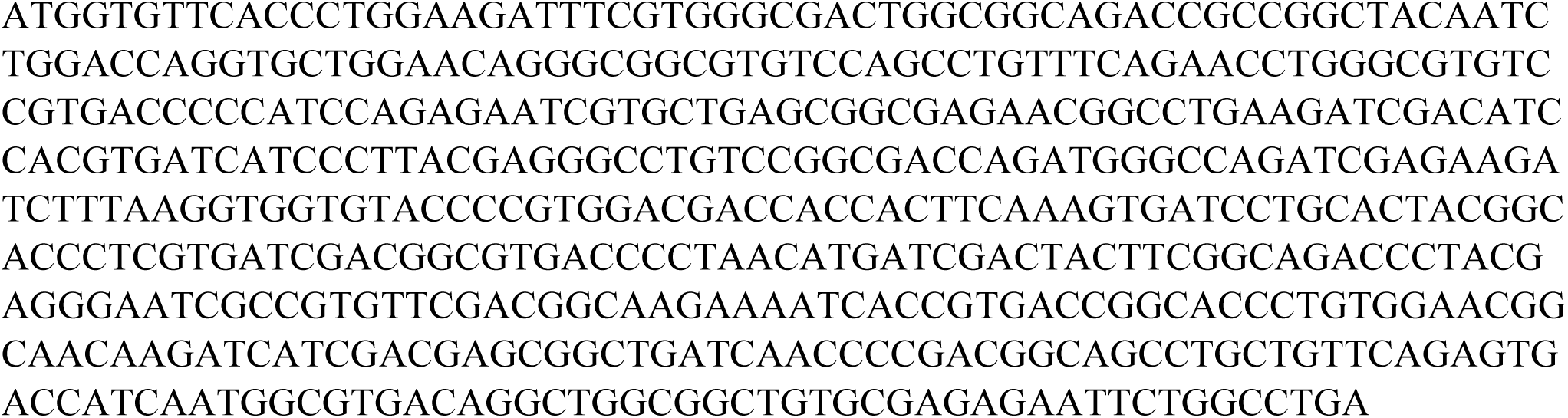
codon optimized NanoLuc NanoLuc codon optimized for human expression using Geneart, changing 87 bp in total.

